# A rapid and dynamic role for FMRP in the plasticity of adult neurons

**DOI:** 10.1101/2023.09.01.555985

**Authors:** Daniel G. Gundermann, Seana Lymer, Justin Blau

## Abstract

Fragile X syndrome (FXS) is a neuro-developmental disorder caused by silencing *Fmr1*, which encodes the RNA-binding protein FMRP. Although *Fmr1* is expressed in adult neurons, it has been challenging to separate acute from chronic effects of loss of *Fmr1* in models of FXS. We have used the precision of *Drosophila* genetics to test if *Fmr1* acutely affects adult neuronal plasticity *in vivo*, focusing on the s-LNv circadian pacemaker neurons that show 24 hour rhythms in structural plasticity. We found that over-expressing *Fmr1* for only 4 hours blocks the activity-dependent expansion of s-LNv projections without altering the circadian clock or activity-regulated gene expression. Conversely, acutely reducing *Fmr1* expression prevented s-LNv projections from retracting. One FMRP target that we identified in s-LNvs is *sif*, which encodes a Rac1 GEF. Our data indicate that FMRP normally reduces *sif* mRNA translation at dusk to reduce Rac1 activity. Overall, our data reveal a previously unappreciated rapid and direct role for FMRP in acutely regulating neuronal plasticity in adult neurons, and underscore the importance of RNA-binding proteins in this process.

## Introduction

Cytoplasmic RNA-binding proteins (RBPs) regulate gene expression post-transcriptionally by controlling mRNA localization, stability and translation. One advantage of regulating gene expression via RBPs is the ability to rapidly transition between different states of cells. This is probably most clear after fertilization when a large set of maternally-deposited mRNAs are rapidly activated for translation^1^. RBPs are also very important in neurons, where they help localize mRNAs to distal parts of the cell^2^ and control translation locally in response to changes in neuronal activity^3^. These two functions of RBPs are key for neuronal plasticity, which underlies learning and memory^3^ and can be disrupted in various neurological disorders^4^.

Fragile X Syndrome (FXS) exemplifies the importance of RBPs in neurons. FXS is a neuro- developmental disorder which is the most common cause of intellectual disability in humans and also the most common single-gene cause of autism^5–7^. FXS is caused by expansion of CGG repeats in the 5’ UTR of the *Fragile X messenger ribonucleoprotein 1* (*Fmr1*) gene on the X chromosome that silences *Fmr1* expression^8,9^. *Fmr1* encodes an RNA-binding protein known as FMRP, whose canonical function is to inhibit translation of specific mRNAs^10^. However, FMRP can also increase mRNA translation^11^, regulate splicing^12^ and even physically interact with ion channels to regulate neuronal excitability^13,14^.

*Fmr1* is expressed in many tissues, but is enriched in the brain, where its expression is developmentally regulated in mice^15^ and *Drosophila*^16^. For example, the highest FMRP levels in mouse somatosensory cortex coincide with the critical period of sensory-dependent plasticity, and *Fmr1* mutant mice have abnormal structural and synaptic plasticity during this time^17,18^.

*Fmr1* is also expressed in adult neurons^15,16,19^ and plasticity phenotypes in the hippocampus of *Fmr1* mutant mice can be rescued by reintroducing *Fmr1* in adult mice^20^, suggesting a role for FMRP in adult neurons. Indeed, FMRP is localized to dendritic spines^21,22^, which are key sites of neuronal plasticity. However, it is challenging to test acute functions of FMRP *in vivo* in mice.

We chose to use *Drosophila* to explore an acute role of FMRP in adult neuronal plasticity because of the tools available to acutely alter gene expression in adult neurons. *Drosophila* mutant for *Fmr1* show developmental abnormalities such as reduced axonal pruning of mushroom body gamma neurons during development^16^, which can be considered analogous to the neuro-developmental phenotypes in mice FXS models. *Drosophila Fmr1* is also important in young adult flies where it is required to remove the PDF-Tri neurons^23^. In this study, we focused on the small ventral lateral neurons (s-LNvs) to be able to study a single homogenous cell type and thus avoid the potential confound of diverse FMRP targets in different cells. The s-LNvs are 4 of the ∼75 pacemaker neurons in each brain hemisphere that control circadian rhythms in adult flies^24^. s-LNvs have well-documented and predictable plasticity in the structure of their projections^25^ – described below.

Each s-LNv expresses a set of clock genes including *period* (*per*) and *vrille* (*vri)*^26^. These clock genes form an intracellular molecular clock that generates rhythms in the expression of some clock genes (such as *per* and *vri*) and also in the expression of clock-controlled genes that are often cell-type specific^27^. For example, *Irk1* is an s-LNv clock-controlled gene which encodes an inward rectifier potassium channel that helps control the 24hr rhythms in s-LNv excitability that peak around dawn to promote the morning bout of locomotor activity^27,28^. s-LNvs also show 24hr rhythms in the structure of their projections, which are maximally expanded around dawn and retracted by dusk^25^. This structural plasticity involves making and breaking synaptic connections with a 24hr rhythm^29^ to likely control behavioral outputs. The predictability of these structural changes make s-LNvs ideal to study the molecular mechanisms of plasticity.

Inducing firing in s-LNvs at dusk is sufficient to fully expand s-LNv projections within 2-3 hours^30,31^. Conversely, expressing a mammalian Inward rectifier K+ channel prevents the normal expansion of s-LNvs at dawn^32^. Thus s-LNv excitability seems to drive s-LNv structural plasticity, and this is consistent with the requirement for Mef2 – a transcription factor involved in activity-dependent transcription in many neurons^33^ – in the rhythms of s-LNv structural plasticity^30^. Linking transcriptional changes to the s-LNv cytoskeleton, we previously showed that *Puratrophin-like* (*Pura*) encodes a Rho1 guanine nucleotide exchange factor (GEF) whose rhythmic transcription in s-LNvs peaks at dusk^31^. Rhythmic *Pura* expression generates rhythms in Rho1 activity that retract s-LNv projections at dusk by increasing myosin phosphorylation^31^. However, it is not yet clear what genes promote expansion of s-LNv projections at dawn.

s-LNv projections in *Drosophila Fmr1* mutant flies have expanded projections despite relatively normal molecular clock oscillations^34–36^. Conversely, over-expressing *Fmr1* collapses s-LNv projections^36^, indicating that *Fmr1* can regulate s-LNv plasticity bi-directionally. However, these experiments were performed before s-LNv plasticity had been described and were thus only analyzed at a single time of day. Here we report that changing *Fmr1* expression specifically in adult s-LNvs changes s-LNv projections after only 4 hours of altered *Fmr1* levels. We used RNA interference (RNAi) transgenes to show that *Fmr1* is required for s-LNv projections to retract at dusk. Conversely, we found that overexpressing *Fmr1* in s-LNvs is sufficient to block expansion of s-LNv projections at dawn. We then sought to identify how FMRP controls s-LNv plasticity.

We found that acutely altering *Fmr1* expression does not affect the s-LNv molecular clock or a reporter gene that reports neuronal activity in s-LNvs. Thus *Fmr1* likely acts downstream of the clock and neuronal excitability to control s-LNv plasticity. We used the TRIBE-seq method^37^ to identify FMRP targets in s-LNvs. One FMRP target gene we identified is *sif*, which encodes a GEF for the Rac1 GTPase and which has previously been implicated in regulating synaptic growth at the *Drosophila* neuromuscular junction^38^. The mammalian Sif ortholog – TIAM1 – stabilizes dendrites and synapses in the developing hippocampus^39^, and is required for structural plasticity in neurons in the spinal dorsal horn in a mouse neuropathic pain model^40^. Altered *Tiam1* expression is also associated with major depressive disorder^41^.

We found that *sif* and *Rac1* itself are required for s-LNv projections to expand at dawn, and that transiently over-expressing either *sif* or a constitutively active *Rac1* is sufficient to rapidly expand s-LNv projections at dusk when they are normally retracted. Furthermore, the expanded s-LNv projections observed at dusk when *Fmr1* expression is knocked down can be rescued by simultaneously reducing *sif* expression. Thus we propose that FMRP normally represses *sif* mRNA translation at dusk to ensure that the Pura / Rho1 complex is the dominant small GTPase and that s-LNv projections retract. s-LNv structural plasticity is thus a balance of GTPase activity: *Pura* – regulated transcriptionally – activates Rho1 to retract projections at dusk; while *sif* – regulated post-transcriptionally by *Fmr1* – activates Rac1 to expand s-LNv projections at dawn. Given the extensive similarities between *Drosophila* and mammalian FMRP, we speculate that mammalian FMRP can also act rapidly to control the plasticity of adult mammalian neurons. Overall, our data underscore the importance of RNA-binding proteins in acutely regulating adult neuronal plasticity.

## Results

### FMRP dynamically regulates the daily structural plasticity of s-LNvs

To test if FMRP can dynamically regulate the structural plasticity of adult s-LNvs, we used an inducible system to either overexpress or knockdown *Fmr1* in the LNvs of adult flies. *Pdf* encodes a neuropeptide that is restricted to LNvs in the mature adult brain and so we used the *Pdf-Gal4* driver to restrict transgene expression to LNvs^24^. We also used a temperature-sensitive version of Gal80 – a repressor of Gal4 – that is ubiquitously expressed via the *tubulin* enhancer (*tub-Gal80^ts^*)^42^. This Gal4-Gal80^TS^ combination can temporally control expression of UAS transgenes by shifting the temperature from 19°C to 30°C to inactivate Gal80^TS^ and allow Gal4 to activate transcription. We used UAS transgenes with either an *Fmr1* cDNA (*UAS-Fmr1*) for overexpression, or one of two independent short-hairpin RNAs that target *Fmr1* (*UAS-shFmr1*) to knock down *Fmr1*. The flies also had a *Pdf-RFP* transgene in which the *Pdf* regulatory region expresses a cytosolic Red Fluorescent Protein (RFP)^27^, which fills the LNv projections to visualize and quantify their structure.

Flies developed at 19°C and adult flies were then entrained in Light:Dark (LD) cycles at 19°C. The temperature in the incubator was raised to 30°C to induce either over-expression of *Fmr1* or *shFmr1* expression. ZT (Zeitgeber time) indicates time in a 12 hour : 12 hour LD cycle – with lights on from ZT0 to ZT12, and lights off from ZT12 to ZT24. One temperature shift started at ZT22 – 2 hours before lights on – with brains dissected and fixed 4 hours later at ZT2 (“dawn”), when s-LNv projections are normally fully expanded^25^. The opposite temperature shift started at ZT10 with brains dissected 4 hours later at ZT14 (“dusk”), when s-LNv projections are normally retracted^25^. We used antibodies to mRFP1 to visualize s-LNv projections, and quantified their spread in 3D using a previously developed script^31^.

The data in Figure 1A show that control *Pdf-Gal4, tub-Gal80^ts^, Pdf-RFP* flies without a UAS transgene that were kept at 19°C and then shifted to 30°C for the last 4 hours have the typical expanded s-LNv projections at ZT2 and retracted projections at ZT14. We previously saw the same expanded s-LNv projections at ZT2 and retracted projections at ZT14 when *Pdf-Gal4, tub- Gal80^ts^* control flies expressed a control fluorescent protein via a UAS transgene^31^. We also previously showed that the *Pdf-Gal4, tub-Gal80^ts^* combination does not allow leaky transgene expression even when flies were at 25°C for 12 hours^31^.

**Figure 1.**
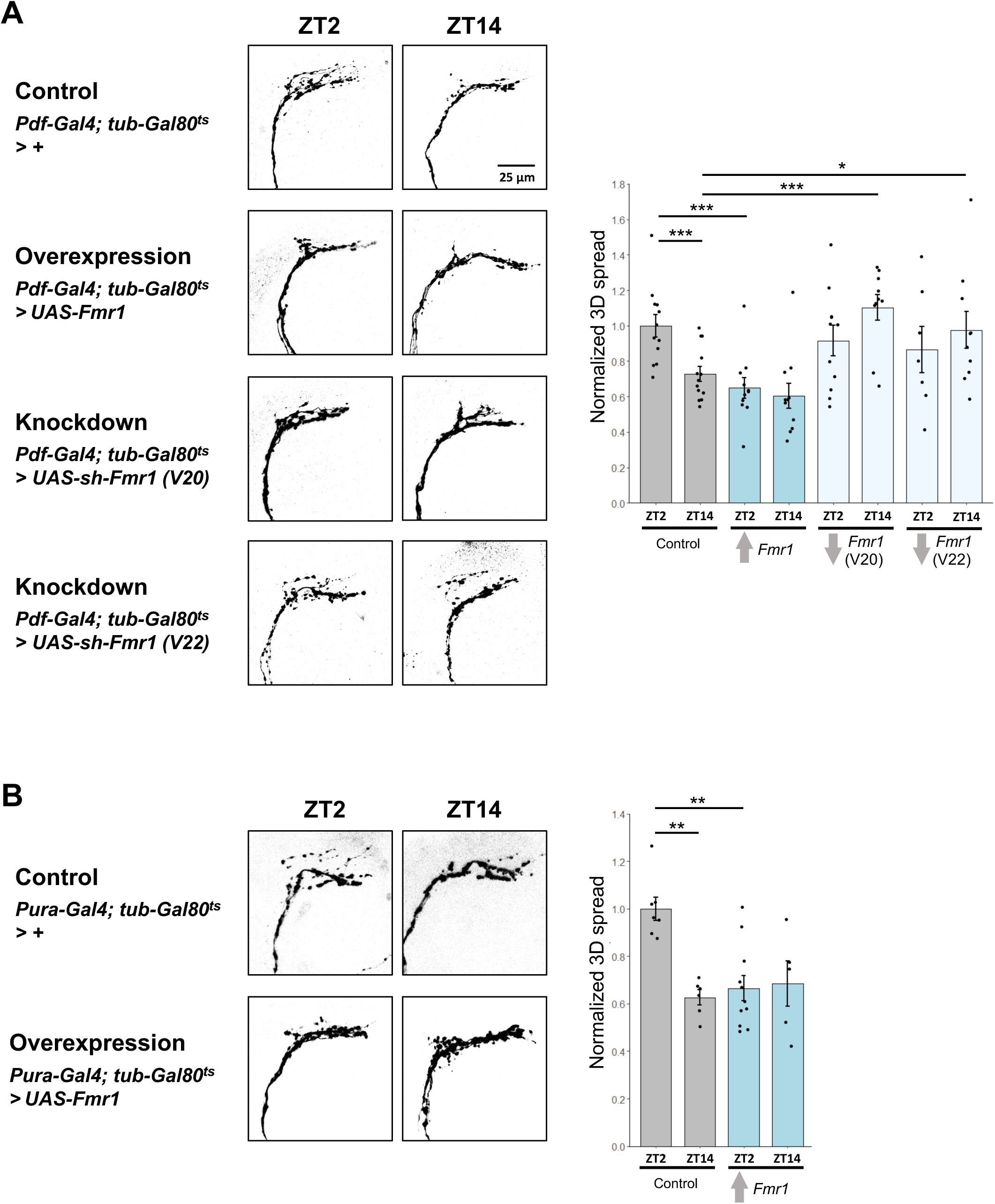
FMRP dynamically regulates the structural plasticity of s-LNvs. (A) Left: Representative confocal images of s-LNv projections from flies with *Pdf-Gal4* and *tubulin-Gal80^ts^* crossed to *w* flies (Control: *Pdf, tub-Gal80^ts^* > *+*), flies with a *UAS-Fmr1* transgene (*Fmr1* overexpression: *Pdf, tub-Gal80^ts^* > *Fmr1*) or to flies with one of two different short hairpin RNA transgenes against *Fmr1*: *Fmr1* knock-down (*Pdf, tub-Gal80^ts^* > *sh-Fmr1 (V20)* and *Pdf, tub-Gal80^ts^* > *sh-Fmr1 (V22*)). Transgenes were induced by shifting flies from 19°C to 30°C for 4 hours and dissecting at either ZT2 or ZT14. All flies also contained a *Pdf-RFP* transgene, and projections were visualized using an antibody to RFP. These images are representative of at least 7 hemispheres from 3 independent experiments. Right: Quantification of the 3D spread of s-LNv projections using a MATLAB script^31^. Bars show the average 3D spread normalized to control s-LNvs at ZT2. Each dot is the 3D spread of one set of s-LNv projections. Error bars show SEM. Significance is denoted by asterisks between the conditions joined by a line. A Wilcoxon test with the adjusted p-value for the number of comparisons of interest was used. * indicates p < 0.05; ** p < 0.01 and *** p<0.001. (B) Left: Representative confocal images as in A of s-LNv projections from Control (*Pura, tub- Gal80^ts^* > *+)*, or *Fmr1* overexpression (*Pura, tub-Gal80^ts^* > *UAS-Fmr1)* flies at either ZT2 or ZT14 after 4 hours of transgene induction. These images are representative of at least 5 hemispheres from 2 independent experiments. Right: Quantification of the 3D spread of the projections as in A. An ANOVA followed by a Tukey post-hoc test was used to assess significance. * indicates p < 0.05; ** p < 0.01 and *** p<0.001.

We found that inducing *Fmr1* overexpression by moving flies to 30°C for 4 hours starting at ZT22 kept s-LNv projections retracted at ZT2. However, we did not observe any difference between control s-LNvs and s-LNvs in which *Fmr1* was overexpressed from ZT10 to ZT14 (Figure 1A). This indicates that over-expressing FMRP is not non-specifically shrinking s-LNv projections – rather FMRP must be blocking a key molecule required for s-LNv projections to expand at dawn.

The data in Figure 1A also show that s-LNv projections stay expanded when either of two different *shFmr1* transgenes were expressed from ZT10 to ZT14 to reduce *Fmr1* expression. However, there was no difference between sLNv projections from control flies and expression of either *shFmr1* transgene from ZT22 to ZT2. The two *Fmr1 shRNA* transgenes are independent as they target different regions of *Fmr1* mRNA and are in two different plasmid backbones. The very similar effects of these two shRNA transgenes allow us to conclude that FMRP is required for s-LNv projections to fully retract at dusk.

The *Pdf-Gal4* driver is expressed in s-LNvs and the large LNvs (l-LNvs), with the latter cells more important for sleep than circadian rhythms^43,44^. To more definitively test if FMRP acts cell-autonomously in s-LNvs to regulate their structural plasticity, we used a different Gal4 driver which expresses in s-LNvs but not in l-LNvs^45^. This driver – *VT027163-Gal4* – has part of the *Pura* regulatory region and we refer to this driver as *Pura-Gal4* from here on. We used *Pura- Gal4* in conjunction with *tub-Gal80^ts^* to induce *Fmr1* overexpression for 4 hours starting at either ZT10 or ZT22 and imaged s-LNv projections 4 hours later as described above. The data in Figure 1B show the normal rhythm in the 3D spread of s-LNv projections from control flies with *Pura-Gal4* and *tub-Gal80^ts^* but no UAS transgene. In contrast, there is no difference in the 3D spread of the s-LNv projections between ZT2 and ZT14 in the experimental flies with *Fmr1* overexpressed, with s-LNv projections retracted at both timepoints.

The data in Figure 1 are consistent with previous analyses of flies carrying *Fmr1* mutations or flies over-expressing *Fmr1*^34–36^. However, our much briefer manipulations allow us to conclude that FMRP can act very rapidly in adult neurons – in this case, s-LNvs – to dynamically regulate their structural plasticity.

### *Fmr1* does not regulate s-LNv structural plasticity through the molecular clock or excitability

The molecular clock in s-LNvs regulates the timing of s-LNv neuronal activity, making them most active around dawn^28^ when s-LNv projections are expanded. In addition, inducing s-LNvs to fire at dusk is sufficient to expand s-LNv projections^30,31^. This leads to a model where the s-LNv molecular clock regulates rhythms in firing, which in turn regulates structural plasticity in an activity-dependent manner. Thus, FMRP could regulate s-LNv structural plasticity by altering the molecular clock in s-LNvs or by altering the timing of s-LNv neuronal activity. These ideas seemed plausible because constitutive long-term overexpression of *Drosophila Fmr1* lengthens circadian period^34^, and because mouse FMRP binds the mRNA of several clock genes including *mPer1* in hippocampal neurons^46^. In addition, mRNAs that regulate neuronal excitability are bound by FMRP, and neuronal excitability is altered in *Fmr1* knockout mice^10,47^.

We first tested if altering *Fmr1* levels for 4 hours changes the phase of the molecular clock by measuring levels of the circadian transcription factor Vri. We chose Vri because its levels are rhythmic in clock neurons, and because its relatively short half-life makes it a more accurate marker of clock phase than longer-lived clock proteins such as Per^48^. We assayed Vri protein levels in s-LNvs by immunofluorescence and quantified the fluorescence intensity at ZT2 and ZT14 after 4 hours of inducing either *Fmr1* over-expression or an *shFmr1* transgene.

Figure 2A shows that control flies have low Vri levels in s-LNvs at ZT2 and high levels at ZT14 as expected^48^. We found no significant difference in Vri levels between control s-LNvs and either experimental group. Since s-LNv structural plasticity is altered before we could detect changes in the s-LNv molecular clock, we conclude that the target(s) that explain the rapid effects of FMRP on s-LNv plasticity are downstream of the molecular clock.

**Figure 2.**
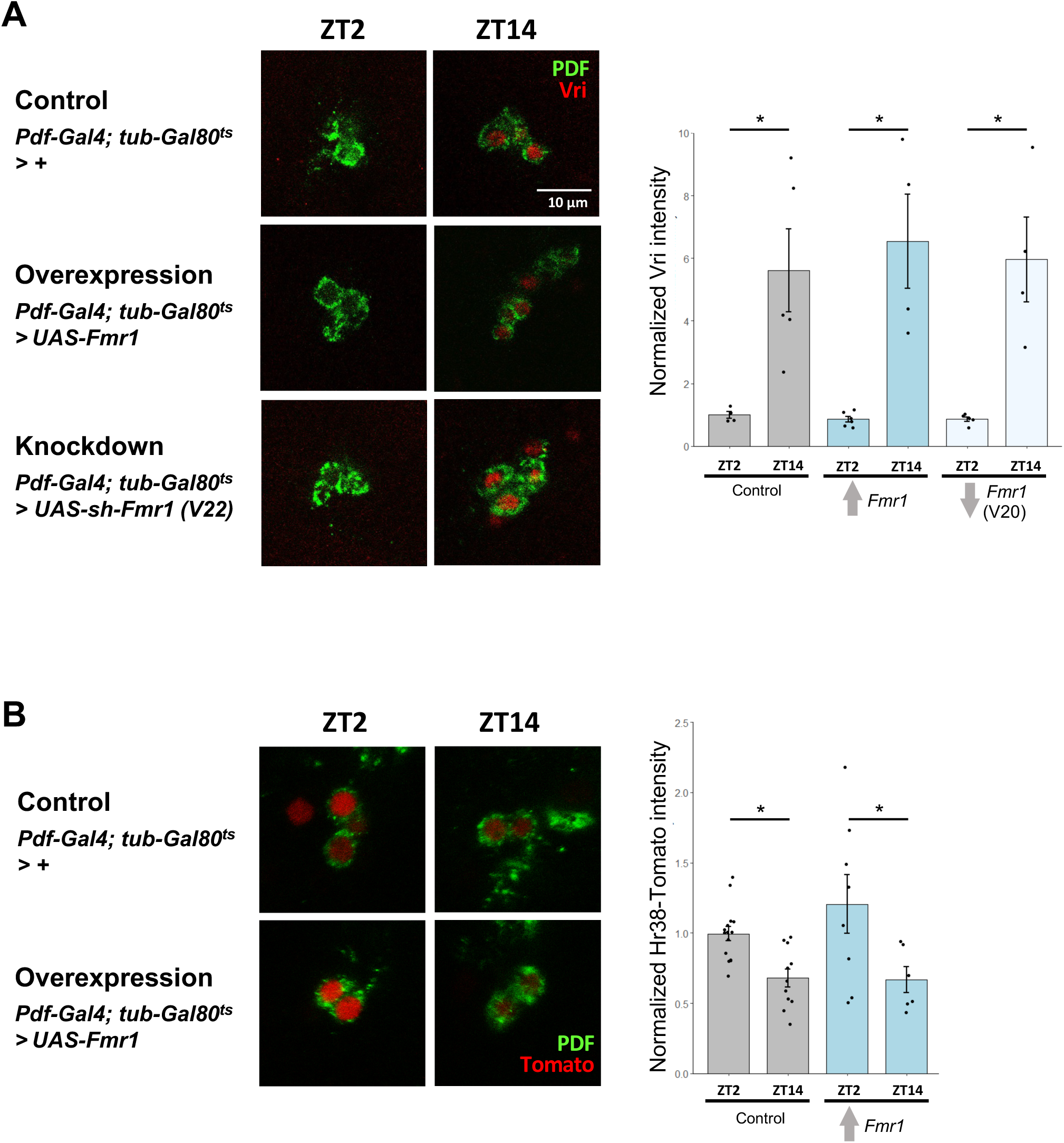
Transiently altering *Fmr1* levels does not affect the s-LNv molecular clock or expression of an activity-dependent reporter gene. (A) Left: Representative images of s-LNv cell bodies from Control (*Pdf, tub-Gal80^ts^* > *+*), *Fmr1* overexpression (*Pdf, tub-Gal80^ts^* > *Fmr1*) or *Fmr1* knock-down flies [*Pdf, tub-Gal80^ts^* > *sh-Fmr1 (V22)]*. Transgenes were induced for 4 hours and brains were dissected at either ZT2 or ZT14. An antibody against Vri (red) was used to measure the molecular clock and s-LNv cell bodies were visualized with an antibody against PDF (green). Right: Quantification of the intensity of Vri signal normalized to levels in control s-LNvs at ZT2 for each genotype. A t-test was used to test for differences between the two times for each genotype. Error bars show SEM. * indicates p < 0.05; ** p < 0.01 and *** p<0.001. These images are representative of at least 4 hemispheres from 3 independent experiments. (B) Left: Representative images of s-LNv cell bodies from Control flies (*Pdf, tub-Gal80^ts^* > *+*) or flies over-expressing *Fmr1* (*Pdf, tub-Gal80^ts^* > *Fmr1*). These flies also have the *Hr38-Tomato* transcriptional reporter gene. Transgenes were induced for 4 hours and brains were dissected at either ZT2 or ZT14, before staining with antibodies to Tomato (red) and PDF (green). Right: Quantification of the intensity of the *Hr38-Tomato* reporter normalized against the levels in control s-LNvs at ZT2. A t-test was used to test for differences between the two times for each genotype. These images are representative of at least 6 hemispheres from 4 independent experiments.

Next we wanted to test if transiently altering *Fmr1* levels rapidly changes the excitability of s-LNvs. Neuronal activity rapidly induces expression of activity-regulated genes – also known as immediate early genes – such as c-fos, Egr1 (also known as NGFI-A and Zif-268) and Nr4a1 (also known as NGFI-B and Nur77)^33^. Expression of activity-regulated genes can be used to determine the recent activity of a neuron *in vivo* (e.g.^49^). While c-fos does not seem to be activity-regulated in insects, the *Drosophila* orthologs of Egr1 – *stripe* (*sr*) – and Nr4a1 – *Hr38* – are upregulated in response to neuronal activity in many different types of neurons^50^, and *Hr38* is also activity-regulated in silkmoths^51^.

We used a transcriptional reporter gene for *Hr38*, in which 4kb of *Hr38* regulatory DNA is upstream of a destabilized and nuclear-localized fluorescent Tomato reporter gene. This *Hr38- Tomato* reporter gene shows higher expression in larval LNvs at ZT2 than at ZT14^52^, consistent with increased larval LNv excitability at dawn^53^, just like adult s-LNvs^28^. Figure 2B shows that control s-LNvs have higher *Hr38-Tomato* levels at ZT2 than at ZT14, and this is not significantly affected by inducing overexpression of *Fmr1* for 4 hours. The similar levels of *Hr38-Tomato* expression at ZT2 with and without *Fmr1* overexpressed make it likely that the rapid changes in s-LNv structural plasticity caused by increasing FMRP levels are downstream of neuronal excitability.

### Identifying FMRP targets *in vivo* in s-LNvs

We wanted to take an unbiased approach to identifying the relevant FMRP target(s) in s-LNvs that mediate the changes in s-LNv plasticity. We decided that it was important to identify targets in s-LNvs themselves since FMRP interactions with other proteins can affect the mRNAs bound by FMRP^54–56^. However, there are only 8 s-LNvs in each *Drosophila* brain out of a total of ∼200,000 neurons^57^, making the task quite challenging. To understand how FMRP can rapidly regulate neuronal plasticity, we also wanted to identify FMRP target mRNAs that interact with FMRP in the timeframe in which we detect s-LNv plasticity phenotypes i.e. within ∼4 hours of altered *Fmr1* expression (Figure 1).

Given these limitations, we decided to use TRIBE-seq^37^. TRIBE stands for **t**argets of **R**NA- binding proteins **i**dentified **b**y **e**diting and uses the enzyme ADAR, which changes adenosines into inosines in mRNA. Inosine is then recognized as a guanosine by polymerases *in vitro* during RNA-sequencing library preparation. Thus the sequence of mRNAs edited by ADAR differs from the genomic sequence. In TRIBE-seq, the ADAR catalytic domain (ADARcd) is fused to an RBP and then expressed in the relevant cells. mRNA is then isolated from these cells and RNA-sequencing used to identify mRNAs with edited adenosines (Fig. 3A). These edited mRNAs must have been physically close to the RBP-ADARcd fusion protein, and thus likely bound by that RBP. McMahon et al^37^ showed that expressing an *Fmr1-ADARcd* transgene in a subset of *Drosophila* excitatory neurons lead to edits in almost half of the orthologs of mammalian FMRP targets identified by cross-linking and immunoprecipitation (CLIP) from mouse brain. These FMRP-ADARcd targets included *futsch*, which had previously been identified genetically and biochemically as a *Drosophila* FMRP target^58^. Furthermore, MAP1B, the mammalian ortholog of *Drosophila futsch*, is one of the top FMRP targets identified by CLIP in the mouse brain^10^.

**Figure 3.**
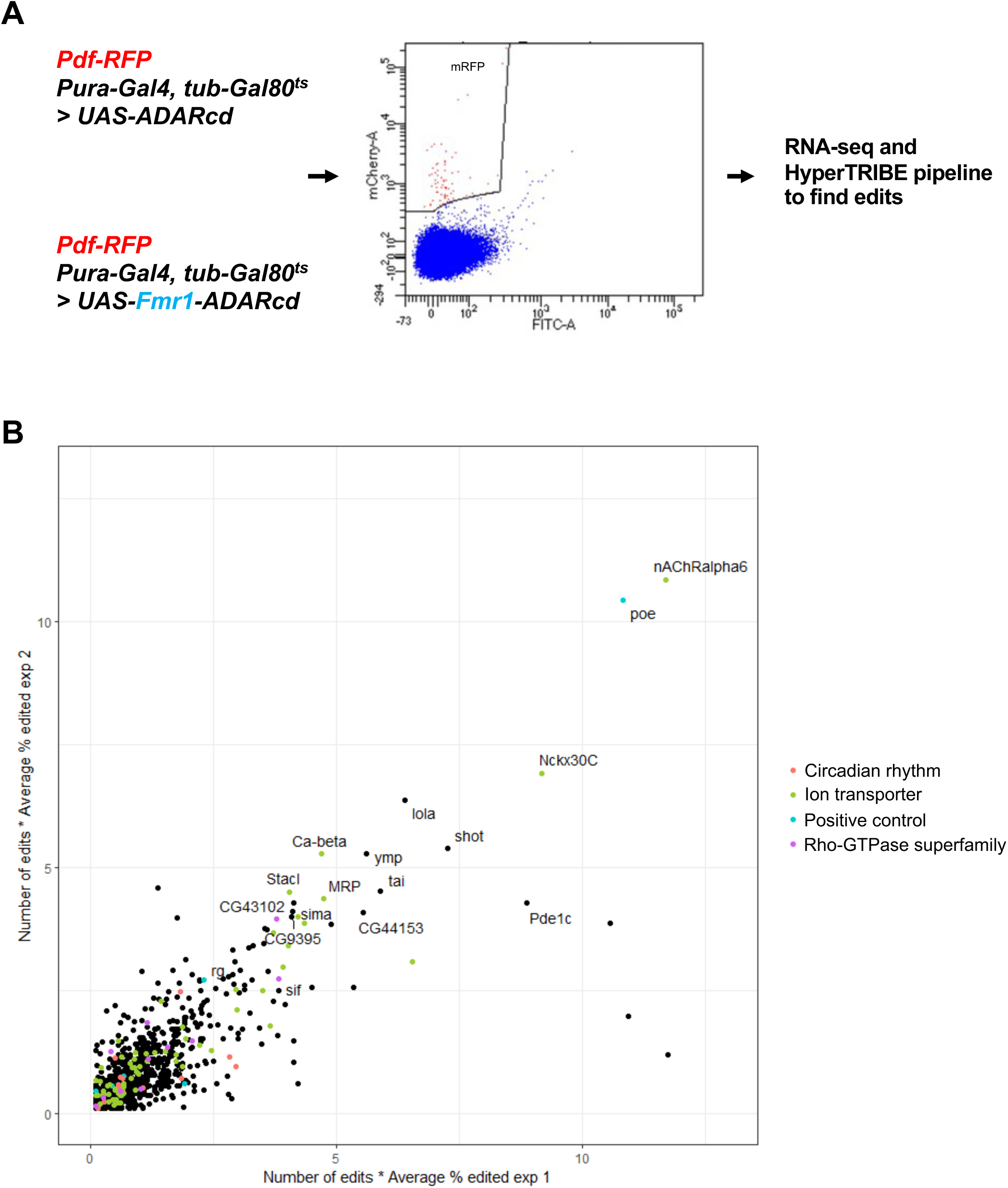
*Fmr1* TRIBE-seq in s-LNvs. (A) Experimental procedure used for *Fmr1* TRIBE-seq in s-LNvs. Genotypes of flies used are shown on the left, followed by an example of FACS sorting of RFP^+^ events from experimental flies. RNA-seq was performed and the HyperTRIBE pipeline^59^ used to identify FMRP targets. (B) Edits from the *Fmr1* TRIBE-seq experiment from experiment 1 and experiment 2 plotted against each other and normalized using the average percentage of edited reads per transcript. Transcripts were color-coded if their GO term is ‘Circadian rhythm’ (GO:0007623, red) or ‘Ion transport’ (GO:0006811, green). A select group of previously identified and validated mammalian and fly FMRP targets are labeled in cyan, and Rho-GTPase superfamily members are in purple.

We used UAS transgenes that express either an FMRP-ADARcd fusion (*UAS-Fmr1-ADARcd*), or only the catalytic domain of a hyperactive version of the ADARcd as a control (*UAS- ADARcd,*^59^). Transgene expression was controlled by *Pura-Gal4* to spatially limit expression to s-LNvs. Temporal control was achieved via the *tubGal80^ts^*transgene, with flies raised and entrained at 19°C to prevent UAS transgene expression. The temperature was then raised to 30°C starting at ZT22 and fly brain dissection began 4 hours later at ZT2. To collect ∼40 brains for each sample, dissections continued for 1 hour, which means that s-LNvs had a range of transgene expression time of between 4 and 5 hours. These flies also contained the constitutively expressed *Pdf-RFP* transgene to ensure a strong enough fluorescent signal for Fluorescence-activated cell sorting (FACS). *Pdf-RFP* expresses in both s-LNvs and l-LNvs, and thus both cell types were purified in this experiment. However, ADAR-modified mRNAs should only come from s-LNvs because *Pura-Gal4* does not express in l-LNvs^45^.

Adult brains were dissociated into a single cell suspension and sorted for RFP+ fluorescence. s-LNvs and particularly l-LNvs have long projections, which sometimes fragment during cell dissociation. Given the localization of RBPs like FMRP outside the cell body^21,22^, and the importance of local translation in neurons^3^, we sorted all RFP+ events via FACS (Figure 3A), having ensured that the collection gates were set to exclude any RFP- cells or cell fragments. We extracted RNA and then generated full-length mRNAs using the SMART-seq3 protocol^60^, followed by RNA-seq as in^37^. We performed two independent experiments with 39-54 brains dissected per condition. The aligned reads were run through the HyperTRIBE pipeline to identify mRNAs modified by FMRP-ADARcd using these previously described cutoff^59^: (i) ≥ 10% of the mRNA reads need to be edited in that location; (ii) ≥ 20 reads for that site; and (iii) no changes to G in the control dataset. The graph in Figure 3B shows the number of edits for a transcript weighted by the average percentage of edits on that transcript. For example, a mRNA with 1 site edited in 60% of reads, and a second site edited in 40% of reads would have a score of 1 (see Methods). We then plotted the weighted edits for the two independent experiments against each other.

We first checked if the relatively brief induction of *Fmr1-ADAR* in s-LNvs was sufficient to identify previously reported targets of *Drosophila* and mammalian FMRP identified by CLIP. We found several FMRP targets including *purity of essence* (*poe* – UBR4 in mammals)^11^, *rugose* (*rg* – Neurobeachin in mammals)^61^, *Calcium/calmodulin-dependent protein kinase II* (*CaMKII*)^62^ and *futsch*^58^(MAP1B in mammals) as FMRP targets in s-LNvs (Figure 3B, all shown in cyan).

Previously described FMRP mRNA targets in neurons include clock genes such as *Per1*^63^, and genes that control neuronal excitability^47,64^. We did not detect many clock genes in our dataset (red in Figure 3B), although some clock genes such as *per* and *vri* are lowly expressed at ZT2 in s-LNvs. We did detect a set of genes encoding ion channels as FMRP targets in s-LNvs (green in Figure 3B). However, these genes are unlikely to explain how rapidly increasing FMRP levels blocks expansion of s-LNv projections given the lack of an effect on the *Hr38-Tomato* activity-dependent transcriptional reporter gene in 4 hours (Figure 2B). We therefore examined the list of FMRP targets for alternative ways to regulate s-LNv plasticity.

We noticed two Rho-GTPase activity regulators in the top 50 mRNAs edited by FMR1-ADAR in s-LNvs: *still life* (*sif*) and *CG43102* (purple in Figure 3B). The presence of these two genes is interesting because expression of the Rho1 GEF *Puratrophin-like* (*Pura*) peaks at dusk in s-LNvs to drive rhythmic Rho1 activity and retract s-LNv projections^31^. The importance of GEFs in s-LNv plasticity was further supported by Polcownuk et al^65^ who showed that over-expressing the GEF *trio* keeps s-LNvs in a dusk-like retracted state even at dawn. [*Pura* itself was not detected in our FMRP TRIBE-seq experiments, although it is not highly expressed at this time of day.] *sif* was also one of the top 20 FMRP targets identified in *Drosophila* cholinergic neurons^37^ and *CG43102* was also identified by FMRP TRIBE-seq^37^, although with many fewer edits than *sif*. Given the importance of Rho GTPases in s-LNv plasticity, we decided to focus on *sif* and *CG43102*.

We also sequenced the genome of flies that contained the genotypes that differed between the two lines used for TRIBE-seq and used the GATK HaplotypeCaller to identify SNPs that could be detected as edits and used the Broad Institute GATK Best Practices recommendations for filtering SNPs. We removed all the A to G SNPs as well as the reverse complementary SNPs that were identified by this analysis from the HyperTRIBE results. Figure S1 shows that *sif* is still in the top targets even after this very stringent cutoff.

### *sif* and *Rac1* regulate the expansion of s-LNv projections at dawn

We first tested whether *sif* and *CG43102* are involved in the daily structural plasticity of s-LNvs. We used the same transient expression strategy as in Figure 1 using *UAS-shRNA* transgenes to reduce expression and either a *UAS-sif* transgene to over-express *sif* or a P-element insertion line (EY01540) that contains UAS binding sites inserted just upstream of *CG43102*.

We found no changes in the structure of s-LNv projections with either strategy for altering *CG43102* expression (Figure S2). However, we cannot formally exclude a role for *CG43102* in s-LNv plasticity as we did not check the efficacy of overexpression or knockdown in the lines we used.

In contrast, we found strong s-LNv plasticity phenotypes from altering *sif* expression. First, we found that over-expressing *sif* for 4 hours from ZT10 to ZT14 leaves s-LNv projections in an expanded state at ZT14, when they are normally retracted. *sif* over-expression from ZT22 to ZT2 had no effect and s-LNv projections remained fully expanded, indicating that the *sif-* mediated expansion of s-LNv projections at ZT14 is a time-specific expansion of s-LNv projections (Figure 4A).

**Figure 4.**
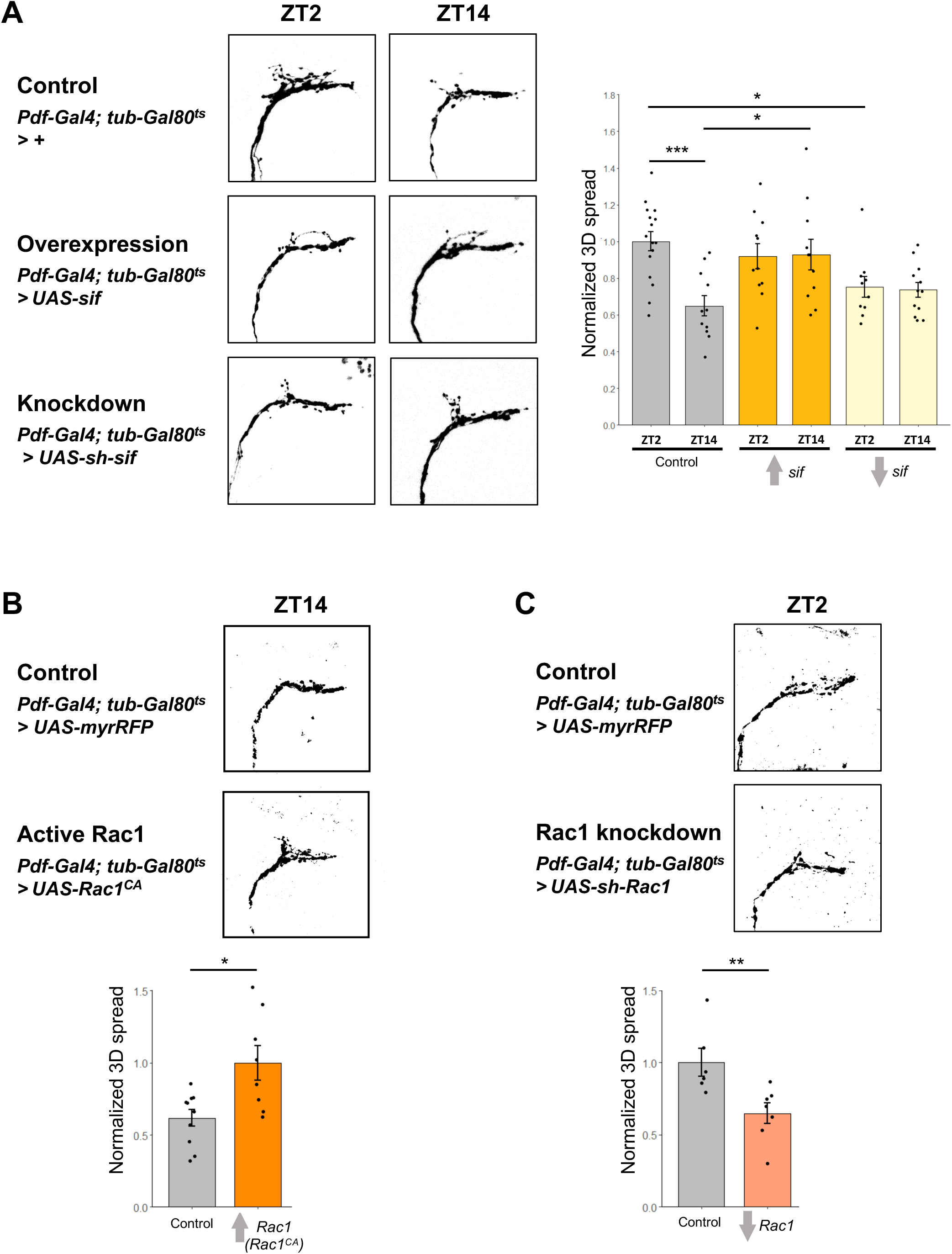
*sif* and *Rac1* regulate the expansion of s-LNv projections at dawn. (A) Left: representative confocal images of s-LNv projections from flies with *Pdf-Gal4* and *tubulin-Gal80^ts^* crossed to *w* flies (Control: *Pdf, tub-Gal80^ts^* > *+*), to flies with a *UAS-sif* transgene (Overexpression: *Pdf, tub-Gal80^ts^* > *UAS-sif*), or to flies with a short hairpin RNA against *sif* (Knockdown: *Pdf, tub-Gal80^ts^* > *UAS-sh-sif).* Transgenes were induced for 4 hours as in Figure 1 and dissected at either ZT2 or ZT14. All flies also contained a *Pdf-RFP* transgene, and projections were visualized using an antibody to RFP. These images are representative of at least 10 hemispheres from 4 independent experiments. Right: Quantification of the 3D spread of s-LNv projections as in Figure 1. An ANOVA followed by a Tukey post-hoc test was used to assess significance. * indicates p < 0.05; ** p < 0.01 and *** p<0.001. (B) Left / Top: Representative confocal images of s-LNv projections from flies with *Pdf-Gal4* and *tubulin-Gal80^ts^* crossed to flies with a *UAS-myrRFP* transgene (Control: *Pdf, tub-Gal80^ts^ > UAS- myrRFP*), or to flies with a constitutively active *UAS-Rac1* transgene (Active Rac1: *Pdf, tub- Gal80^ts^* > *UAS-Rac1^CA^*). These transgenes were induced at ZT12 and dissected 2 hours later at ZT14. s-LNv projections were visualized using an antibody to PDF. Right / Bottom: Quantification of the 3D spread of s-LNv projections as in Figure 1. A t-test was used to test for differences between the two genotypes. Error bars show SEM. * indicates p < 0.05; ** p < 0.01 and *** p<0.001. These images are representative of at least 8 hemispheres from 2 independent experiments. (C) Left / Top: Representative confocal images of s-LNv projections from flies with *Pdf-Gal4* and *tubulin-Gal80^ts^* crossed to flies with a *UAS-myrRFP* transgene (Control: *Pdf, tub-Gal80^ts^ > UAS- myrRFP*), or to flies with a short hairpin RNA targeting *Rac1* (Knockdown: *Pdf, tub-Gal80^ts^* > *UAS-sh-Rac1).* These transgenes were induced at ZT12 and dissected 14 hours later at ZT2. s- LNv projections were visualized using an antibody to PDF. Right / Bottom: Quantification of the 3D spread of s-LNv projections as in Figure 1. A t-test was used to test for differences between the two genotypes. Error bars show SEM. * indicates p < 0.05; ** p < 0.01 and *** p<0.001. These images are representative of at least 6 hemispheres from 2 independent experiments.

In the complementary experiment, we found that inducing expression of *sif-shRNA* from ZT22 to ZT2 lead to retracted s-LNv projections at dawn, in contrast to the expanded s-LNv projections in control flies. However, expressing *sif-shRNA* from ZT10 to ZT14 had no effect, and s-LNv projections stayed retracted. Therefore we conclude that *sif* is normally required for s-LNv projections to expand at dawn. Taking these data together with *sif* being sufficient to expand s-LNv projections when over-expressed at dusk (Figure 4A), we conclude that control of Sif protein levels in s-LNvs is very important for rhythmic structural plasticity in in s-LNvs.

*sif* encodes a Rac1 GEF which localizes to the larval neuromuscular junction and to the synaptic terminal in adult photoreceptor cells^38^. TIAM1 (T cell lymphoma invasion and metastasis 1) is the mammalian Sif ortholog and activates Rac1 after NMDA receptor stimulation to reorganize the actin cytoskeleton in mammalian hippocampal neurons^66^. Interestingly, *sif* was one of only two genes that showed rhythms in its alternative splicing in all three different classes of clock neurons assayed – LN_v_s, DN1s and LNds^67^, further supporting the idea that regulation of *sif* is important for clock neuron function.

Since *sif* encodes a Rac1-GEF, we wanted to test if Rac1 itself can regulate the plasticity of s-LNv projections. We first used a UAS transgene expressing a constitutively active version of Rac1 (*UAS-Rac1^CA^*) that acts on downstream targets independently of GEF activity^68^. We used *Pdf-Gal4* and *tub-Gal80^ts^*to induce expression of *UAS-Rac1^CA^* at dusk, and quantified the 3D spread of s-LNv projections this time using antibodies to PDF. We found that inducing expression of Rac1^CA^ for just 2 hours from ZT12 to ZT14 increased the 3D spread of the s-LNvs when control s-LNv projections are normally retracted (Figure 4B). Conversely, when an RNAi targeting *Rac1* was expressed for 14 hours starting at ZT12 and ending at ZT2, we found that s-LNv projections are retracted (Figure 4C). Thus our data show that Rac1 is normally involved in expanding s-LNv projections at dawn, and that uncoupling the regulation of Rac1 from a GEF is sufficient to rapidly expand s-LNv projections when they are normally retracted.

### A genetic interaction between *sif* and *Fmr1*

We have shown that s*if* mRNA is an FMRP target in s-LNvs (Figure 3B), that *sif* is required at dawn for s-LNv projections to expand, and that abnormally high *sif* mRNA levels at dusk keep s-LNv projections expanded (Figure 4A). We therefore hypothesized that FMRP represses *sif* mRNA translation at dusk to help retract s-LNv projections.

We used genetics to test this idea by asking if the abnormally expanded s-LNv projections at dusk caused by *Fmr1-shRNA* can be rescued by co-expressing *sif-shRNA*. Our logic was as follows: If FMRP normally represses *sif* translation at dusk, then FMRP knockdown at dusk increases *sif* mRNA translation, leading to expanded s-LNv projections. Thus co-expressing *sif- shRNA* alongside *Fmr1-shRNA* might return s-LNv projections to their normally retracted ZT14 phenotype if the FMRP-*sif* mRNA interaction is important for s-LNv structural plasticity.

We again induced expression of the transgenes for 4 hours using *Pdf-Gal4* and *tub-Gal80^ts^*. The results in Figure 5 confirm that expressing *Fmr1-shRNA* alone from ZT10 to ZT14 prevented s-LNv projections from retracting, as in Figure 1. However, we found that co-expressing *Fmr1- shRNA* with *sif-shRNA* gave retracted s-LNv projections at ZT14 that were not significantly different from control s-LNvs at ZT14. Note that expressing *sif-shRNA* alone did not significantly affect the 3D spread of s-LNv projections (see Figure 4A). To control for the possibility that expressing a second shRNA transgene diminishes the effect of the *UAS-Fmr1-shRNA* transgene, we used the *UAS-CG43102-shRNA* transgene as a control. We found that co-expressing *Fmr1-shRNA* and *CG43102-shRNA* gave expanded projections that were not significantly different from expressing *Fmr1-shRNA* alone. Thus, the rescue of the *Fmr1-shRNA* phenotype with *sif-shRNA* is not a trivial consequence of overwhelming the RNAi pathway or the Gal4/UAS system by expressing 2 UAS-shRNA transgenes. Instead, we conclude that reducing *sif* expression specifically overrides the defect caused by reduced *Fmr1* expression. This result is consistent with FMRP normally repressing *sif* translation at dusk to allow s-LNvs to retract.

**Figure 5.**
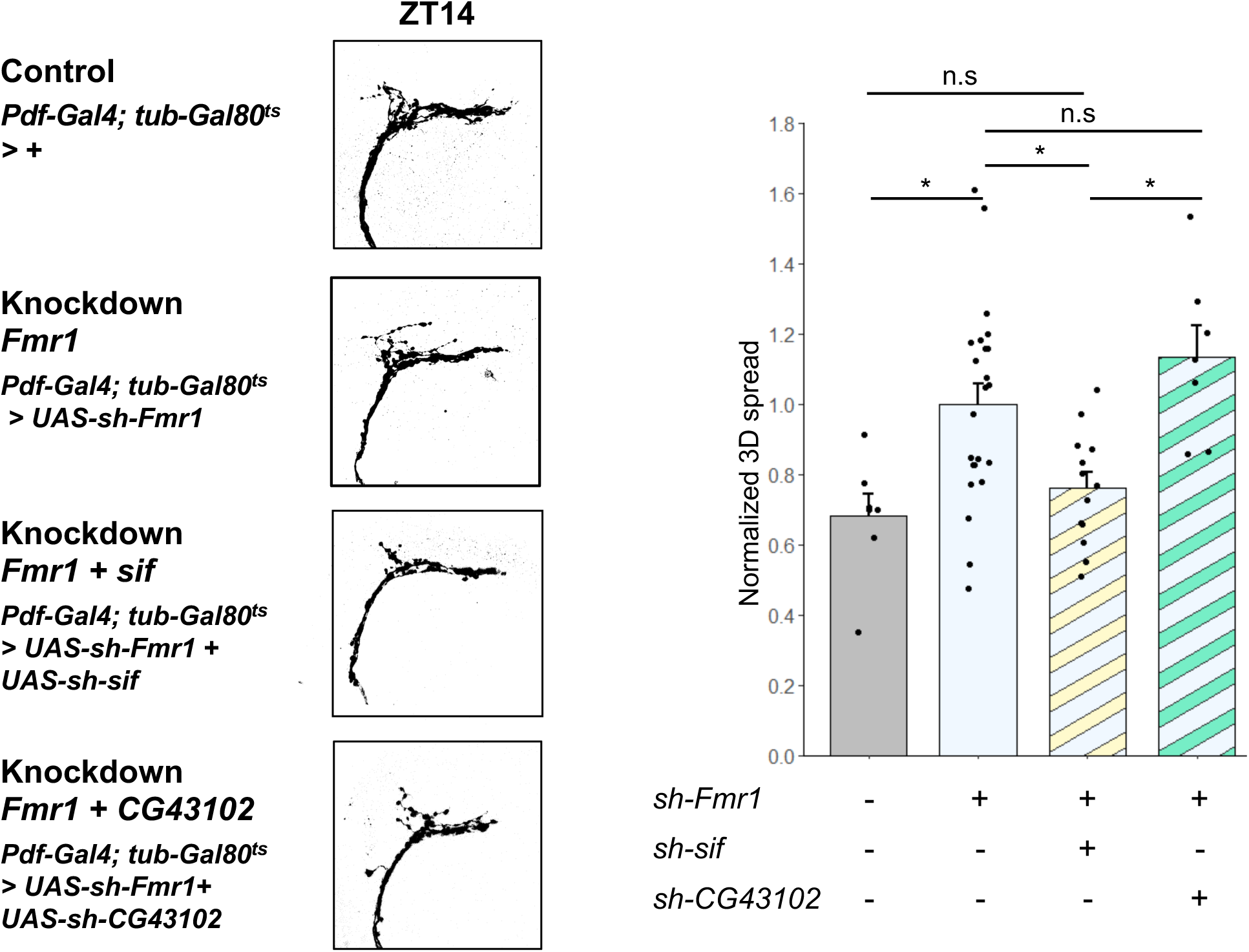
*sif* knockdown can rescue the effect on the s-LNv projections of the *Fmr1* knockdown. Left: Representative confocal images of s-LNv projections from flies with *Pdf-Gal4* and *tubulin- Gal80^ts^* crossed to *w* flies (Control: *Pdf, tub-Gal80^ts^* > *+*), flies with an shRNA transgene against *Fmr1* only (*Fmr1* knockdown: *Pdf, tub-Gal80^ts^* > *shFmr1*), flies with shRNA transgenes against both *Fmr1* and *sif* (*Fmr1* + *sif* knock-down: *Pdf, tub-Gal80^ts^* > *sh-Fmr1 + sh-sif)* or flies with shRNA transgenes against both *Fmr1* and *CG43102* (*Fmr1* + *CG43102* knock-down: *Pdf, tub- Gal80^ts^* > *sh-Fmr1* + *sh-CG43102*). Transgenes were induced by moving flies to 30°C at ZT10 hours and dissecting 4 hours later at ZT14. All flies also contained a *Pdf-RFP* transgene, and projections were visualized using an antibody to RFP. These images are representative of at least 7 hemispheres from 6 independent experiments. Right: Quantification of the 3D spread of s-LNv projections as in Figure 1. An ANOVA followed by a Tukey post-hoc test was used to assess significance. * indicates p < 0.05; ** p < 0.01 and *** p<0.001.

## Discussion

It has been challenging to address the roles of FMRP in mature neurons given the effects of loss of *Fmr1* on neuronal development. We used *Drosophila* genetics to test the acute role of FMRP in the plasticity of adult neurons and found that FMRP rapidly and bi-directionally regulates the plasticity of adult s-LNv circadian pacemaker neurons: Too much FMRP blocks the activity-dependent expansion of s-LNv projections at dawn; and too little FMRP prevents their retraction at dusk. Working in this single cell type, we identified a set of mRNAs targeted by FMRP and we showed that one target – a Rac1 GEF encoded by *sif* – explains at least some of the s-LNv cellular phenotypes caused by altering *Fmr1* expression.

Our study adds to the roles of RBPs in *Drosophila* circadian pacemaker neurons, where RBPs can regulate circadian period length. One mechanism for this is the regulation of *per* mRNA translation by the RBPs Atx-2 and Tyf^69–71^. Another example is the RBP Lark which targets *double-time* (*dbt*) RNA in clock neurons^72^. *dbt* encodes the kinase that controls PER protein stability and thus circadian period length^73^. However, our study is the first to our knowledge to find a role for an RBP in s-LNv structural plasticity.

Identifying *sif* as an FMRP target in s-LNvs allowed us to establish that Sif / Rac1 is normally involved in expanding s-LNv projections at dawn. Our data support the idea that FMRP dynamically regulates the structural plasticity of s-LNvs by reducing translation of *sif* mRNA at dusk (Figure 6). This should reduce Rac1 activity at the same time as rising *Pura* mRNA levels increase Rho1 activity^31^. It is relatively common for cells to decrease one process simultaneous with increases in the opposing process. This can help turn the output of two individual graded signals into a switch – for example, in the interplay between transcriptional activators and repressors that bind the same or overlapping DNA-sequences^74,75^. An analogous process with small GTPases in s-LNvs could help ensure that their projections are either expanding or retracting, but not trying to do both simultaneously. Indeed, s-LNv plasticity phenotypes tend to leave projections either constitutively expanded or constitutively retracted, and rarely stuck in between states (Figures 1, 4, 5)^30,31^. In addition, crosstalk between Rac1 and Rho1 GTPases themselves has been observed^76^ and this could also help ensure switch-like behavior between expansion and retraction in s-LNv projections. We also note that the pre-synaptic protein Bruchpilot (Brp) associates with Sif and RhoGAP100F^77^. Thus Sif could also be involved in the daily rhythms observed in the number of active zones in s-LNvs projections^29,31^. In addition, it is intriguing that Brp binds RhoGAP100F which should inactivate Rho1 and could thereby help the switch from Rho1 to Rac1 activity, although this remains to be tested. Finally, it should also be noted that FMRP itself indirectly regulates the Rac1-Wave regulatory complex via its interactions with CYFIP1 in mice^78,79^.

**Figure 6.**
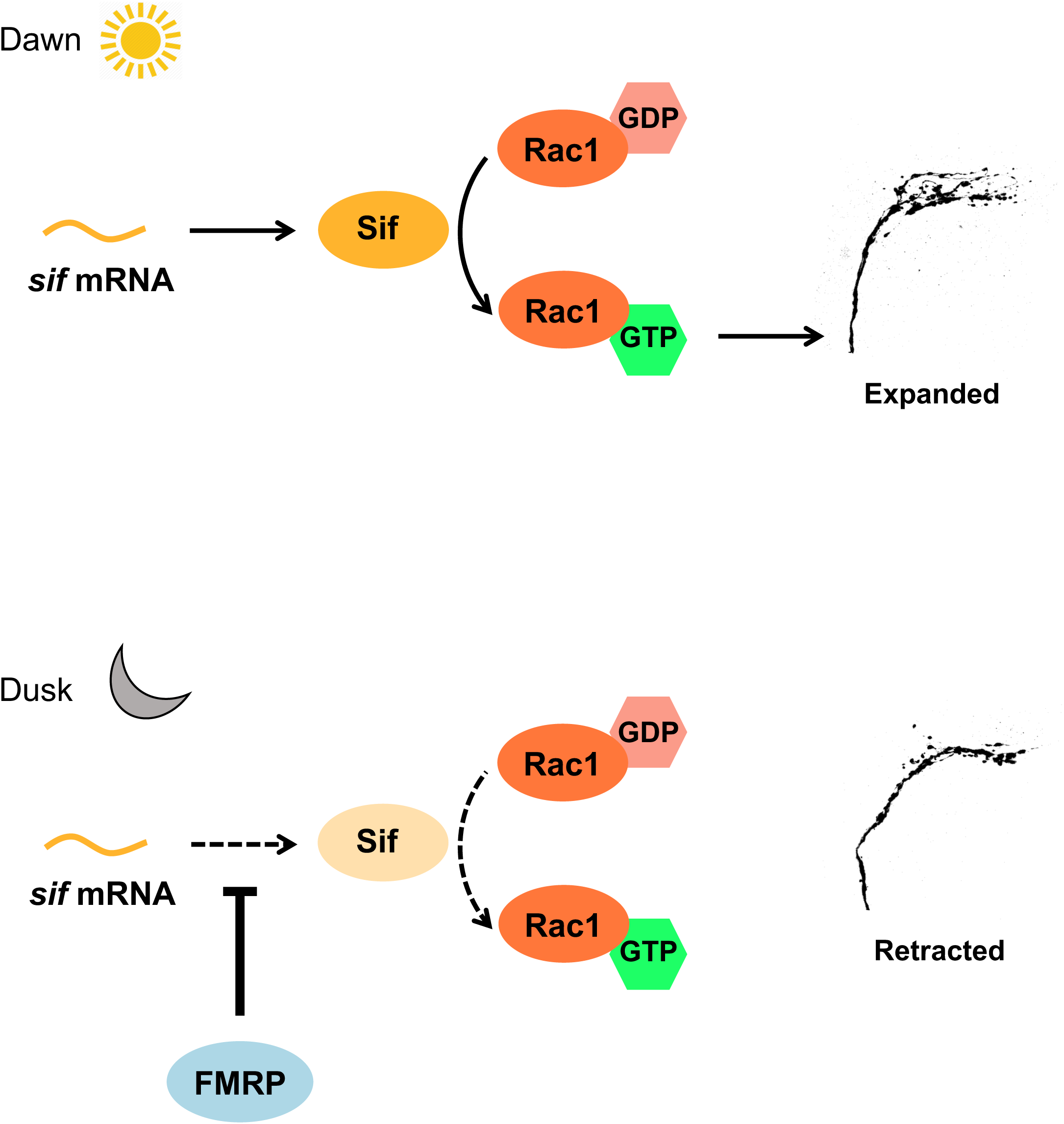
Model to explain how FMRP rapidly controls s-LNv structural plasticity. *sif* mRNA is translated around dawn, which allows Sif protein to activate Rac1, which in turn expands s-LNv projections. An increase in the activity of FMRP around dusk reduces *sif* mRNA translation, thereby reducing activation of Rac1 and helping to retract s-LNv projections.

For FMRP to dynamically regulate Sif protein levels, FMRP activity must change over 24 hours such that FMRP is more active at dusk than dawn. FMRP activity could be regulated at any level from transcription to post-transcriptional modifications. Mammalian FMRP is dephosphorylated in response to neuronal activity, which disaggregates FMRP-mRNA granules, thereby increasing translation of mRNAs that were previously bound by FMRP^80^. Such a mechanism could help explain how experimentally increasing neuronal activity at dusk expands s-LNv projections in only 2 hours^30^. However, phosphorylation of FMRP is unlikely to fully explain how neuronal activity regulates s-LNv plasticity: Over-expressing FMRP at dawn blocks expansion of s-LNv projections even though s-LNvs seem to continue to fire normally as shown by the activity-dependent *Hr38* transcriptional reporter gene (Figure 2B). For additional mechanisms of regulating FMRP activity, we note that *Fmr1* mRNA levels are lower in LNvs that have been chronically hyper-excited^81^, suggesting that *Fmr1* mRNA levels are regulated by neuronal activity. In addition, translation of *Fmr1* mRNA can also be regulated by neuronal activity^21^.

TRIBE-seq also identified *sif* mRNA as an FMRP target in cholinergic and GABAergic neurons in *Drosophila*^37^, raising the possibility that the mechanism described here is a general mechanism for controlling neuronal plasticity. Indeed, Tiam1 was identified as an FMRP target in mouse brains using CLIP^10^. This idea is further supported by the genetic interaction of *Fmr1* and *Rac1* in the development of dendrites in the *Drosophila* peripheral nervous system^82^.

The post-translational regulation of Sif / Rac1 activity described here contrasts with the transcriptional regulation of Pura / Rho1 activity that drives retraction of s-LNv projections at dusk^31^. However, the requirement of Mef2 in expanding s-LNv projections^30^ suggests that activity-dependent transcriptional regulation is also important at dawn. Thus s-LNv structural plasticity is similar to the plasticity associated with learning and memory by using both transcription factors and RNA-binding proteins to regulate gene expression^3,83^, making s-LNvs a good model to study the fundamental molecular mechanisms of neuronal plasticity. In conclusion, our data reveal a previously unappreciated rapid and direct role for FMRP in regulating neuronal plasticity in adult neurons, and underscore the importance of RNA-binding proteins in this process.

## Methods

### Fly rearing and transgenes

For transgene induction experiments, flies were raised at 19°C and then entrained as adults for at least 4 days in 12:12 light-dark cycles at 19°C. To induce transgene expression, the incubator temperature was changed to 30°C for 2, 4 or 14 hours before dissections. Transgenic flies used were as follows, with Bloomington Stock IDs in parentheses:

**Gal4 and Gal80 lines:** *Pdf-Gal4* (80939)^24^; *Pura-Gal4* (VT027163-Gal4), described in^45^; *tub- Gal80^ts^* (7018)^42^.

**Reporter genes:** *Pdf-RFP*^27^; *Hr38-Tomato*^52^

**UAS transgenes:** *UAS-Fmr1* (6931)^58^; *UAS-ADARcd* (UAS-ADAR-E488Q from^84^; *UAS-Fmr1- ADAR*^37^; *UAS-sif* (9127)^85^; *UAS-myrRFP*^86^; *UAS-Rac1^CA^* (6291)^68^.

**UAS shRNA lines:** *UAS-sh-Fmr1 (V20)* (34944); *UAS-sh-Fmr1 (V22)* (35200); *UAS-sh-sif* (61934); *UAS-sh-CG43102* (62371); *UAS-sh-Rac1* (34910), previously shown to reduce Rac1 expression in fly heads when crossed to *elav-Gal4*^87^).

We used the *EY01540* EPgy2 insertion line inserted just upstream of *CG43102* for attempted over-expression (Bloomington Stock: 15515).

### Immunofluorescence and imaging

Immunofluorescence was done similarly to ref.^31^. Briefly, dissected brain samples were fixed in 4% formaldehyde for 20 minutes at room temperature before permeabilizing with 1% Triton X- 100 in PBS and blocking with 10% heat-inactivated goat serum in PBS with 0.5% Triton X-100 (PBST). Samples were incubated with primary antibodies in the same PBST buffer with serum at 4°C for two nights in all experiments except for Figure 4B, which was for one night. After washing in PBST, samples were incubated with the appropriate secondary antibodies in PBST + 10% serum for 1 hour at room temperature before further washing in PBST. Finally, samples were allowed to sink in 50% glycerol and then mounted on a glass slide and coverslip with SlowFade™ Gold Antifade Mountant (ThermoFisher). Slides were stored at 4°C in the dark until imaging on a Leica SP8 confocal microscope for all images except those in Figure 4B, which were taken on a Leica SP5. Brains were imaged on the SP8 with the 63x lens at 1.94x zoom and a resolution of 1024x1024. Antibodies used were Takara Living Colors dsRed polyclonal rabbit antibody at 1:500 to recognize RFP, mouse anti-PDF C7 at 1:50^88^ (provided by the Developmental Studies Hybridoma Bank), and guinea Pig anti-Vri at 1:5000, generously provided by Paul Hardin.

### Quantification of the 3D spread of s-LNv projections

s-LNv projections were quantified using a custom MATLAB script as described previously^31^. Briefly, the confocal stacks were saved as separate JPEG files using ImageJ. The script was then run on those images and a polygon drawn around the dorsal projections starting where the projections turn. The script generates the variation along the x, y and z axes, and these values are multiplied together to determine the spread in 3 dimensions (3D spread) of the projections. The 3D spread data were then loaded into RStudio and each condition tested for normality with the Shapiro-Wilk method. This normality test was only significant for Figure 1A. for which a Kruskal-Wallis test followed by a Wilcoxon test was used between the conditions of interest, and the p-value was adjusted using the ‘p.adjust’ function in R with the Benjamini-Hochberg method with 7 as the number of comparisons. For all other figures, an ANOVA was performed followed by a Tukey HSD post-hoc test to test for significant differences between the conditions.

### Quantification of Vri and Hr38-Tomato levels

ImageJ was used to quantify the intensity in s-LNv cell nuclei. The intensity value was obtained by drawing a polygon around the desired area and then using the ‘Measure’ function in ImageJ. An area outside the cell bodies was recorded as background, and the average of the s-LNv intensity across 4 cells was divided by the background to get a single datapoint.

### FMRP TRIBE-seq in s-LNvs

Control flies had the genotype: *Pdf-RFP* / *UAS-ADARcd*; *Pura-Gal4, tub-Gal80^ts^* / *Sb* or *TM6B* Experimental flies had the genotype: *Pdf-RFP* / *UAS-Fmr1-ADARcd*; *Pura-Gal4, tub-Gal80^ts^* / *+*. Flies were raised at 19°C and then transgene expression was induced at ZT22 by shifting to 30°C. Brains were dissected in Schneider’s media (add company – Sigma?) for 1 hour from ZT1.5 to ZT2.5 and then dissociation was started by incubating a 0.5% Trypsin solution in PBS for 30 minutes at 25°C in a shaking dry bath. Brains were then washed with ice-cold Schneider’s media and then with ice-cold D-PBS before pipetting up and down 50 times for mechanical disassociation. Tissue clumps were removed by passing the sample through a 20µm filter.

RFP+ events were sorted using a BD FACS Aria II (BD Biosciences). A control sample with no RFP transgene was used to set a gate to exclude non-fluorescent events. All samples were kept on ice while they waited to be sorted. RFP+ fragments were sorted into 300μL of Dynabeads™ mRNA Purification Kit lysis buffer. Samples were frozen at -80°C until processing.

A Dynabeads™ mRNA purification kit was used to extract mRNA. SMART-seq3^60^ was performed for bulk sorted events using 15 amplification cycles. Libraries were prepared with the Illumina XT DNA kit from 1 ng of amplified cDNA with 6 cycles of amplification while adding indices. Library sizes were checked using a Tapestation (Agilent Technologies) to ensure a peak centered around 300-800bp. Libraries were sequenced using an Illumina NextSeq500 with paired-end reads of 150bp.

### Analysis of TRIBE-seq data

Trimmomatic was used to trim sequencing data with the following parameters:

-*phred33 HEADCROP:6 LEADING:25 TRAILING:25 AVGQUAL:25 MINLEN:19*.

Sequences were aligned to the *Drosophila* genome version dm6 using STAR and the following parameters:

*--outFilterMismatchNoverLmax 0.07 --outFileNamePrefix aligned_ADARcd_exp2 -- outFilterMatchNmin 16 --outFilterMultimapNmax 1*.

Samtools was then used to filter alignments which had a higher than 10 phred score with Samtools view and the *-q 10* parameter. Duplicates were removed using Picard. The HyperTRIBE protocol^59^ was then followed to obtain the edited sites of the *Fmr1-ADARcd* condition using *ADARcd* RNA-seq as the control. The final file of the HyperTRIBE pipeline produced by the script ‘*summarize_results.pl*’ was imported into RStudio for analysis.

For Supplemental Figure 1 genomic DNA was extracted from *UAS-ADARcd; Sb / TM6b* and *UAS-Fmr1-ADAR* flies using standard procedures. Libraries were prepared using the NEBNext Ultra II FS DNA Library Prep Kit for Illumina using 30 minutes of fragmentation in the first step. Libraries were sequenced with MiSeq 2x75 - 150 Cycle v3. Sequences were aligned to the *Drosophila* genome version dm6 using BWA-MEM and SNPs identified using GATK HaplotypeCaller with the parameters recommended by the Broad institute: *QD < 2.0, FS > 60.0, MQ < 40.0, SOR > 4.0, MQRankSum < -12.5* and *ReadPosRankSum < -8.0.* Finally the .vcf files were filtered to find SNPs that change from an A to G or the reverse complement T to C in the ADARcd to the Fmr1-ADARcd condition. Finally these SNPs were converted into a .bed file and subtracted from the .bedgraph files obtained from the HyperTRIBE pipeline using bedtools.

## Acknowledgements

We are very grateful to Michael Rosbash, Kate Abruzzi and Aoife MacMahon for TRIBE flies and for sharing unpublished data. We are also very grateful to Eugenia Olesnicky for suggesting that we use TRIBE-Seq, to Bassem Hassan for first encouraging us to study FMRP many years ago, to Eric Lai for advice throughout the course of this project and to Christine Vogel, David Owald and Tom Jongens for helpful discussions. We thank Eric Klann and Jennifer Lennon for comments on the manuscript. We thank the *Drosophila* Transgenic RNAi Project at Harvard Medical School (NIGMS R01-GM084947) and the Bloomington *Drosophila* Stock Center for flies, and Paul Hardin and the Developmental Studies Hybridoma Bank for antibodies. We also thank Simon Kidd, Sofia Luminari and Angelina Xu for help with dissections. This investigation was conducted in facilities constructed with support from Research Facilities Improvement Grant Number C06 RR-15518-01 from the NIH National Center for Research Resources. DG and SL were partly supported by the NYU’s Graduate School of Arts & Science MacCracken Program. This work was supported by NIH grant GM136363 to JB.

## References

1 Winata, C. L. & Korzh, V. The translational regulation of maternal mRNAs in time and space. FEBS Lett 592, 3007–3023 (2018). 10.1002/1873-3468.13183

2 Abouward, R. & Schiavo, G. Walking the line: mechanisms underlying directional mRNA transport and localisation in neurons and beyond. Cell Mol Life Sci 78, 2665–2681 (2021). 10.1007/s00018-020-03724-3

3 Ule, J. & Darnell, R. B. RNA binding proteins and the regulation of neuronal synaptic plasticity. Curr Opin Neurobiol 16, 102–110 (2006). 10.1016/j.conb.2006.01.003

4 Thelen, M. P. & Kye, M. J. The role of RNA binding proteins for local mRNA translation: Implications in neurological disorders. Front Mol Biosci 6, 161 (2019). 10.3389/fmolb.2019.00161

5 Nelson, D. L., Orr, H. T. & Warren, S. T. The unstable repeats--three evolving faces of neurological disease. Neuron 77, 825–843 (2013). 10.1016/j.neuron.2013.02.022

6 Wang, L. W., Berry-Kravis, E. & Hagerman, R. J. Fragile X: leading the way for targeted treatments in autism. Neurotherapeutics 7, 264–274 (2010). 10.1016/j.nurt.2010.05.005

7 Schaefer, G. B. & Mendelsohn, N. J. Genetics evaluation for the etiologic diagnosis of autism spectrum disorders. Genet Med 10, 4–12 (2008). 10.1097/GIM.0b013e31815efdd7

8 Pieretti, M. et al. Absence of expression of the *FMR-1* gene in Fragile X syndrome. Cell 66, 817–822 (1991). 10.1016/0092-8674(91)90125-i

9 Verkerk, A. J. et al. Identification of a gene (*FMR-1*) containing a CGG repeat coincident with a breakpoint cluster region exhibiting length variation in fragile X syndrome. Cell 65, 905–914 (1991). 10.1016/0092-8674(91)90397-h

10 Darnell, J. C. et al. FMRP stalls ribosomal translocation on mRNAs linked to synaptic function and autism. Cell 146, 247–261 (2011). 10.1016/j.cell.2011.06.013

11 Greenblatt, E. J. & Spradling, A. C. Fragile X mental retardation 1 gene enhances the translation of large autism-related proteins. Science 361, 709–712 (2018). 10.1126/science.aas9963

12 Zhou, L. T. et al. A novel role of Fragile X Mental Retardation Protein in pre-mRNA alternative splicing through RNA-binding protein 14. Neuroscience 349, 64–75 (2017). 10.1016/j.neuroscience.2017.02.044

13 Brown, M. R. et al. Fragile X Mental Retardation Protein controls gating of the sodium-activated potassium channel Slack. Nat Neurosci 13, 819–821 (2010). 10.1038/nn.2563

14 Deng, P. Y. et al. FMRP regulates neurotransmitter release and synaptic information transmission by modulating action potential duration via BK channels. Neuron 77, 696–711 (2013). 10.1016/j.neuron.2012.12.018

15 Gholizadeh, S., Halder, S. K. & Hampson, D. R. Expression of Fragile X Mental Retardation Protein in neurons and glia of the developing and adult mouse brain. Brain Res 1596, 22–30 (2015). 10.1016/j.brainres.2014.11.023

16 Tessier, C. R. & Broadie, K. *Drosophila* Fragile X Mental Retardation Protein developmentally regulates activity-dependent axon pruning. Development 135, 1547–1557 (2008). 10.1242/dev.015867

17 Bureau, I., Shepherd, G. M. & Svoboda, K. Circuit and plasticity defects in the developing somatosensory cortex of *FMR1* knock-out mice. J Neurosci 28, 5178–5188 (2008). 10.1523/JNEUROSCI.1076-08.2008

18 Harlow, E. G. et al. Critical period plasticity is disrupted in the barrel cortex of *FMR1* knockout mice. Neuron 65, 385–398 (2010). 10.1016/j.neuron.2010.01.024

19 Tamanini, F. et al. Differential expression of *FMR1*, *FXR1* and *FXR2* proteins in human brain and testis. Hum Mol Genet 6, 1315–1322 (1997). 10.1093/hmg/6.8.1315

20 Zeier, Z. et al. Fragile X Mental Retardation Protein replacement restores hippocampal synaptic function in a mouse model of fragile X syndrome. Gene Ther 16, 1122–1129 (2009). 10.1038/gt.2009.83

21 Weiler, I. J. et al. Fragile X Mental Retardation Protein is translated near synapses in response to neurotransmitter activation. Proc Natl Acad Sci U S A 94, 5395–5400 (1997). 10.1073/pnas.94.10.5395

22 Ferrari, F. et al. The Fragile X Mental Retardation Protein-RNP granules show an mGluR-dependent localization in the post-synaptic spines. Mol Cell Neurosci 34, 343–354 (2007). 10.1016/j.mcn.2006.11.015

23 Gatto, C. L. & Broadie, K. Fragile X Mental Retardation Protein is required for programmed cell death and clearance of developmentally-transient peptidergic neurons. Dev Biol 356, 291–307 (2011). 10.1016/j.ydbio.2011.05.001

24 Renn, S. C., Park, J. H., Rosbash, M., Hall, J. C. & Taghert, P. H. A *pdf* neuropeptide gene mutation and ablation of PDF neurons each cause severe abnormalities of behavioral circadian rhythms in *Drosophila*. Cell 99, 791–802 (1999). 10.1016/s0092-8674(00)81676-1

25 Fernandez, M. P., Berni, J. & Ceriani, M. F. Circadian remodeling of neuronal circuits involved in rhythmic behavior. PLoS Biol 6, e69 (2008). 10.1371/journal.pbio.0060069

26 Allada, R. & Chung, B. Y. Circadian organization of behavior and physiology in *Drosophila*. Annu Rev Physiol 72, 605–624 (2010). 10.1146/annurev-physiol-021909-135815

27 Ruben, M., Drapeau, M. D., Mizrak, D. & Blau, J. A mechanism for circadian control of pacemaker neuron excitability. J Biol Rhythms 27, 353–364 (2012). 10.1177/0748730412455918

28 Cao, G. & Nitabach, M. N. Circadian control of membrane excitability in *Drosophila melanogaster* lateral ventral clock neurons. J Neurosci 28, 6493–6501 (2008). 10.1523/JNEUROSCI.1503-08.2008

29 Gorostiza, E. A., Depetris-Chauvin, A., Frenkel, L., Pirez, N. & Ceriani, M. F. Circadian pacemaker neurons change synaptic contacts across the day. Curr Biol 24, 2161–2167 (2014). 10.1016/j.cub.2014.07.063

30 Sivachenko, A., Li, Y., Abruzzi, K. C. & Rosbash, M. The transcription factor Mef2 links the *Drosophila* core clock to Fas2, neuronal morphology, and circadian behavior. Neuron 79, 281–292 (2013). 10.1016/j.neuron.2013.05.015

31 Petsakou, A., Sapsis, T. P. & Blau, J. Circadian rhythms in Rho1 activity regulate neuronal plasticity and network hierarchy. Cell 162, 823–835 (2015). 10.1016/j.cell.2015.07.010

32 Depetris-Chauvin, A. et al. Adult-specific electrical silencing of pacemaker neurons uncouples molecular clock from circadian outputs. Curr Biol 21, 1783–1793 (2011). 10.1016/j.cub.2011.09.027

33 Flavell, S. W. & Greenberg, M. E. Signaling mechanisms linking neuronal activity to gene expression and plasticity of the nervous system. Annu Rev Neurosci 31, 563–590 (2008). 10.1146/annurev.neuro.31.060407.125631

34 Dockendorff, T. C. et al. *Drosophila* lacking *dfmr1* activity show defects in circadian output and fail to maintain courtship interest. Neuron 34, 973–984 (2002). 10.1016/s0896-6273(02)00724-9

35 Sekine, T. et al. Circadian phenotypes of *Drosophila* Fragile X mutants in alternative genetic backgrounds. Zoolog Sci 25, 561–571 (2008). 10.2108/zsj.25.561

36 Gatto, C. L. & Broadie, K. Temporal requirements of the Fragile X Mental Retardation Protein in modulating circadian clock circuit synaptic architecture. Front Neural Circuits 3, 8 (2009). 10.3389/neuro.04.008.2009

37 McMahon, A. C. et al. TRIBE: Hijacking an RNA-editing enzyme to identify cell-specific targets of RNA-binding proteins. Cell 165, 742–753 (2016). 10.1016/j.cell.2016.03.007

38 Sone, M. et al. Still life, a protein in synaptic terminals of *Drosophila* homologous to GDP-GTP exchangers. Science 275, 543–547 (1997). 10.1126/science.275.5299.543

39 Cheng, J. et al. The Rac-GEF Tiam1 promotes dendrite and synapse stabilization of dentate granule cells and restricts hippocampal-dependent memory functions. J Neurosci 41, 1191–1206 (2021). 10.1523/JNEUROSCI.3271-17.2020

40 Li, L. et al. Tiam1 coordinates synaptic structural and functional plasticity underpinning the pathophysiology of neuropathic pain. Neuron 111, 2038–2050 e2036 (2023). 10.1016/j.neuron.2023.04.010

41 Aston, C., Jiang, L. & Sokolov, B. P. Transcriptional profiling reveals evidence for signaling and oligodendroglial abnormalities in the temporal cortex from patients with major depressive disorder. Mol Psychiatry 10, 309–322 (2005). 10.1038/sj.mp.4001565

42 McGuire, S. E., Le, P. T., Osborn, A. J., Matsumoto, K. & Davis, R. L. Spatiotemporal rescue of memory dysfunction in *Drosophila*. Science 302, 1765–1768 (2003). 10.1126/science.1089035

43 Parisky, K. M. et al. PDF cells are a GABA-responsive wake-promoting component of the *Drosophila* sleep circuit. Neuron 60, 672–682 (2008). 10.1016/j.neuron.2008.10.042

44 Sheeba, V. et al. Large ventral lateral neurons modulate arousal and sleep in *Drosophila*. Curr Biol 18, 1537–1545 (2008). 10.1016/j.cub.2008.08.033

45 Sekiguchi, M., Inoue, K., Yang, T., Luo, D. G. & Yoshii, T. A catalog of GAL4 drivers for labeling and manipulating circadian clock neurons in *Drosophila melanogaster*. J Biol Rhythms 35, 207–213 (2020). 10.1177/0748730419895154

46 Sawicka, K. et al. FMRP has a cell-type-specific role in CA1 pyramidal neurons to regulate autism-related transcripts and circadian memory. Elife 8 (2019). 10.7554/eLife.46919

47 Contractor, A., Klyachko, V. A. & Portera-Cailliau, C. Altered neuronal and circuit excitability in Fragile X Syndrome. Neuron 87, 699–715 (2015). 10.1016/j.neuron.2015.06.017

48 Cyran, S. A., et al. *vrille*, *Pdp1*, and *dClock* form a second feedback loop in the *Drosophila* circadian clock. Cell 112, 329–341 (2003). 10.1016/s0092-8674(03)00074-6

49 Lin, D. et al. Functional identification of an aggression locus in the mouse hypothalamus. Nature 470, 221–226 (2011). 10.1038/nature09736

50 Chen, X., Rahman, R., Guo, F. & Rosbash, M. Genome-wide identification of neuronal activity-regulated genes in *Drosophila*. Elife 5 (2016). 10.7554/eLife.19942

51 Fujita, N. et al. Visualization of neural activity in insect brains using a conserved immediate early gene, *Hr38*. Curr Biol 23, 2063–2070 (2013). 10.1016/j.cub.2013.08.051

52. Zhu, Z., Sanchez Ortiz, T., Mezan, S., Kadener, S. & Blau, J. Transcription of a plasticity gene is activated by neuronal hyperpolarization. bioRxiv 636878, doi: 10.1101/636878 (2019).

53 Collins, B., Kane, E. A., Reeves, D. C., Akabas, M. H. & Blau, J. Balance of activity between LNvs and glutamatergic dorsal clock neurons promotes robust circadian rhythms in *Drosophila*. Neuron 74, 706–718 (2012). 10.1016/j.neuron.2012.02.034

54 Worpenberg, L. et al. Ythdf is a N6-methyladenosine reader that modulates Fmr1 target mRNA selection and restricts axonal growth in *Drosophila*. EMBO J 40, e104975 (2021). 10.15252/embj.2020104975

55 Bardoni, B., Schenck, A. & Mandel, J. L. A novel RNA-binding nuclear protein that interacts with the Fragile X Mental Retardation (FMR1) protein. Hum Mol Genet 8, 2557–2566 (1999). 10.1093/hmg/8.13.2557

56 Napoli, I. et al. The Fragile X Syndrome protein represses activity-dependent translation through CYFIP1, a new 4E-BP. Cell 134, 1042–1054 (2008). 10.1016/j.cell.2008.07.031

57 Raji, J. I. & Potter, C. J. The number of neurons in *Drosophila* and mosquito brains. PLoS One 16, e0250381 (2021). 10.1371/journal.pone.0250381

58 Zhang, Y. Q. et al. *Drosophila* Fragile X-related gene regulates the MAP1B homolog Futsch to control synaptic structure and function. Cell 107, 591–603 (2001). 10.1016/s0092-8674(01)00589-x

59 Rahman, R., Xu, W., Jin, H. & Rosbash, M. Identification of RNA-binding protein targets with HyperTRIBE. Nat Protoc 13, 1829–1849 (2018). 10.1038/s41596-018-0020-y

60 Hagemann-Jensen, M. et al. Single-cell RNA counting at allele and isoform resolution using Smart-seq3. Nat Biotechnol 38, 708–714 (2020). 10.1038/s41587-020-0497-0

61 Sears, J. C., Choi, W. J. & Broadie, K. Fragile X Mental Retardation Protein positively regulates PKA anchor Rugose and PKA activity to control actin assembly in learning/memory circuitry. Neurobiol Dis 127, 53–64 (2019). 10.1016/j.nbd.2019.02.004

62 Sudhakaran, I. P. et al. FMRP and Ataxin-2 function together in long-term olfactory habituation and neuronal translational control. Proc Natl Acad Sci U S A 111, E99–E108 (2014). 10.1073/pnas.1309543111

63 Tang, X. et al. FMRP binds *Per1* mRNA and downregulates its protein expression in mice. Mol Brain 16, 33 (2023). 10.1186/s13041-023-01023-z

64 Gibson, J. R., Bartley, A. F., Hays, S. A. & Huber, K. M. Imbalance of neocortical excitation and inhibition and altered UP states reflect network hyperexcitability in the mouse model of Fragile X syndrome. J Neurophysiol 100, 2615–2626 (2008). 10.1152/jn.90752.2008

65 Polcownuk, S., Yoshii, T. & Ceriani, M. F. Decapentaplegic acutely defines the connectivity of central pacemaker neurons in *Drosophila*. J Neurosci 41, 8338–8350 (2021). 10.1523/JNEUROSCI.0397-21.2021

66 Tolias, K. F. et al. The Rac1-GEF Tiam1 couples the NMDA receptor to the activity-dependent development of dendritic arbors and spines. Neuron 45, 525–538 (2005). 10.1016/j.neuron.2005.01.024

67 Wang, Q., Abruzzi, K. C., Rosbash, M. & Rio, D. C. Striking circadian neuron diversity and cycling of *Drosophila* alternative splicing. Elife 7 (2018). 10.7554/eLife.35618

68 Luo, L., Liao, Y. J., Jan, L. Y. & Jan, Y. N. Distinct morphogenetic functions of similar small GTPases: *Drosophila* Drac1 is involved in axonal outgrowth and myoblast fusion. Genes Dev 8, 1787–1802 (1994). 10.1101/gad.8.15.1787

69 Lim, C. et al. The novel gene twenty-four defines a critical translational step in the *Drosophila* clock. Nature 470, 399–403 (2011). 10.1038/nature09728

70 Lim, C. & Allada, R. ATAXIN-2 activates PERIOD translation to sustain circadian rhythms in *Drosophila*. Science 340, 875–879 (2013). 10.1126/science.1234785

71 Zhang, Y., Ling, J., Yuan, C., Dubruille, R. & Emery, P. A role for *Drosophila* ATX2 in activation of PER translation and circadian behavior. Science 340, 879–882 (2013). 10.1126/science.1234746

72 Huang, Y., McNeil, G. P. & Jackson, F. R. Translational regulation of the DOUBLETIME/CKIdelta/epsilon kinase by LARK contributes to circadian period modulation. PLoS Genet 10, e1004536 (2014). 10.1371/journal.pgen.1004536

73 Price, J. L. et al. double-time is a novel *Drosophila* clock gene that regulates PERIOD protein accumulation. Cell 94, 83–95 (1998). 10.1016/s0092-8674(00)81224-6

74 Stanojevic, D., Small, S. & Levine, M. Regulation of a segmentation stripe by overlapping activators and repressors in the *Drosophila* embryo. Science 254, 1385–1387 (1991). 10.1126/science.1683715

75 Rossi, F. M., Kringstein, A. M., Spicher, A., Guicherit, O. M. & Blau, H. M. Transcriptional control: rheostat converted to on/off switch. Mol Cell 6, 723–728 (2000). 10.1016/s1097-2765(00)00070-8

76 Guilluy, C., Garcia-Mata, R. & Burridge, K. Rho protein crosstalk: another social network? Trends Cell Biol 21, 718–726 (2011). 10.1016/j.tcb.2011.08.002

77 Owald, D. et al. A Syd-1 homologue regulates pre- and postsynaptic maturation in *Drosophila*. J Cell Biol 188, 565–579 (2010). 10.1083/jcb.200908055

78 De Rubeis, S. et al. CYFIP1 coordinates mRNA translation and cytoskeleton remodeling to ensure proper dendritic spine formation. Neuron 79, 1169–1182 (2013). 10.1016/j.neuron.2013.06.039

79 Santini, E. et al. Reducing eIF4E-eIF4G interactions restores the balance between protein synthesis and actin dynamics in Fragile X syndrome model mice. Sci Signal 10 (2017). 10.1126/scisignal.aan0665

80 Tsang, B. et al. Phosphoregulated FMRP phase separation models activity-dependent translation through bidirectional control of mRNA granule formation. Proc Natl Acad Sci U S A 116, 4218–4227 (2019). 10.1073/pnas.1814385116

81 Mizrak, D. et al. Electrical activity can impose time of day on the circadian transcriptome of pacemaker neurons. Curr Biol 22, 1871–1880 (2012). 10.1016/j.cub.2012.07.070

82 Lee, A. et al. Control of dendritic development by the *Drosophila* Fragile X-related gene involves the small GTPase Rac1. Development 130, 5543–5552 (2003). 10.1242/dev.00792

83 Kandel, E. R. The molecular biology of memory storage: a dialogue between genes and synapses. Science 294, 1030–1038 (2001). 10.1126/science.1067020

84 Xu, W., Rahman, R. & Rosbash, M. Mechanistic implications of enhanced editing by a HyperTRIBE RNA-binding protein. RNA 24, 173–182 (2018). 10.1261/rna.064691.117

85 Sone, M. et al. Synaptic development is controlled in the periactive zones of *Drosophila* synapses. Development 127, 4157–4168 (2000). 10.1242/dev.127.19.4157

86 Chen, Y. et al. Cell-type-specific labeling of synapses in vivo through synaptic tagging with recombination. Neuron 81, 280–293 (2014). 10.1016/j.neuron.2013.12.021

87 Kikuchi, M. et al. Disruption of a RAC1-centred network is associated with Alzheimer’s disease pathology and causes age-dependent neurodegeneration. Hum Mol Genet 29, 817–833 (2020). 10.1093/hmg/ddz320

88 Cyran, S. A., et al. The Double-time protein kinase regulates the subcellular localization of the *Drosophila* clock protein Period. J Neurosci 25, 5430–5437 (2005). 10.1523/JNEUROSCI.0263-05.2005

