## Supplemental figures for "A rapid and dynamic role for FMRP in the plasticity of adult neurons"

**Gundermann et al - supplementary figure legends**

**Figure S1. *Fmr1* TRIBE-seq in s-LNvs with all possible SNPs removed.**

Edits from the *Fmr1* TRIBE-seq experiment from experiment 1 and experiment 2 plotted against each other and normalized using the average percentage of edited reads per transcript. These are the edits left after removing all possible SNP locations found by sequencing genomic DNA from the *UAS-ADARcd; Sb / TM6b* and *UAS-Fmr1-ADAR* flies.

**Figure S2. Manipulating *CG43102* expression does not change s-LNv structural plasticity**

(A) Left: representative confocal images of s-LNv projections from flies with *Pdf-Gal4* and *tubulin-Gal80$^{ts}$* crossed to:

(i) *w* flies (Control: *Pdf, tub-Gal80$^{ts}$* > +);

or (ii) flies with the EY01540 P-element that contains UAS binding sites inserted 100bp upstream of the major start site of *CG43102* transcription (Overexpression: *Pdf, tub-Gal80$^{ts}$* > *CG43102*$^{EY01540}$);

or (iii) flies with a short hairpin RNA against *CG43102* (Knockdown: *Pdf, tub-Gal80$^{ts}$* > *UAS-sh-CG43102).*

Transgenes were induced for 4 hours as in Figure 1 and dissected at either ZT2 or ZT14. All flies also contained a *Pdf-RFP* transgene, and projections were visualized using an antibody to RFP. These images are representative of at least 6 hemispheres from 2 independent experiments.

Right: Quantification of the 3D spread of s-LNv projections as in Figure 1. An ANOVA followed by a Tukey post-hoc test was used to assess significance. * indicates $p < 0.05$; ** $p < 0.01$ and *** $p<0.001$.

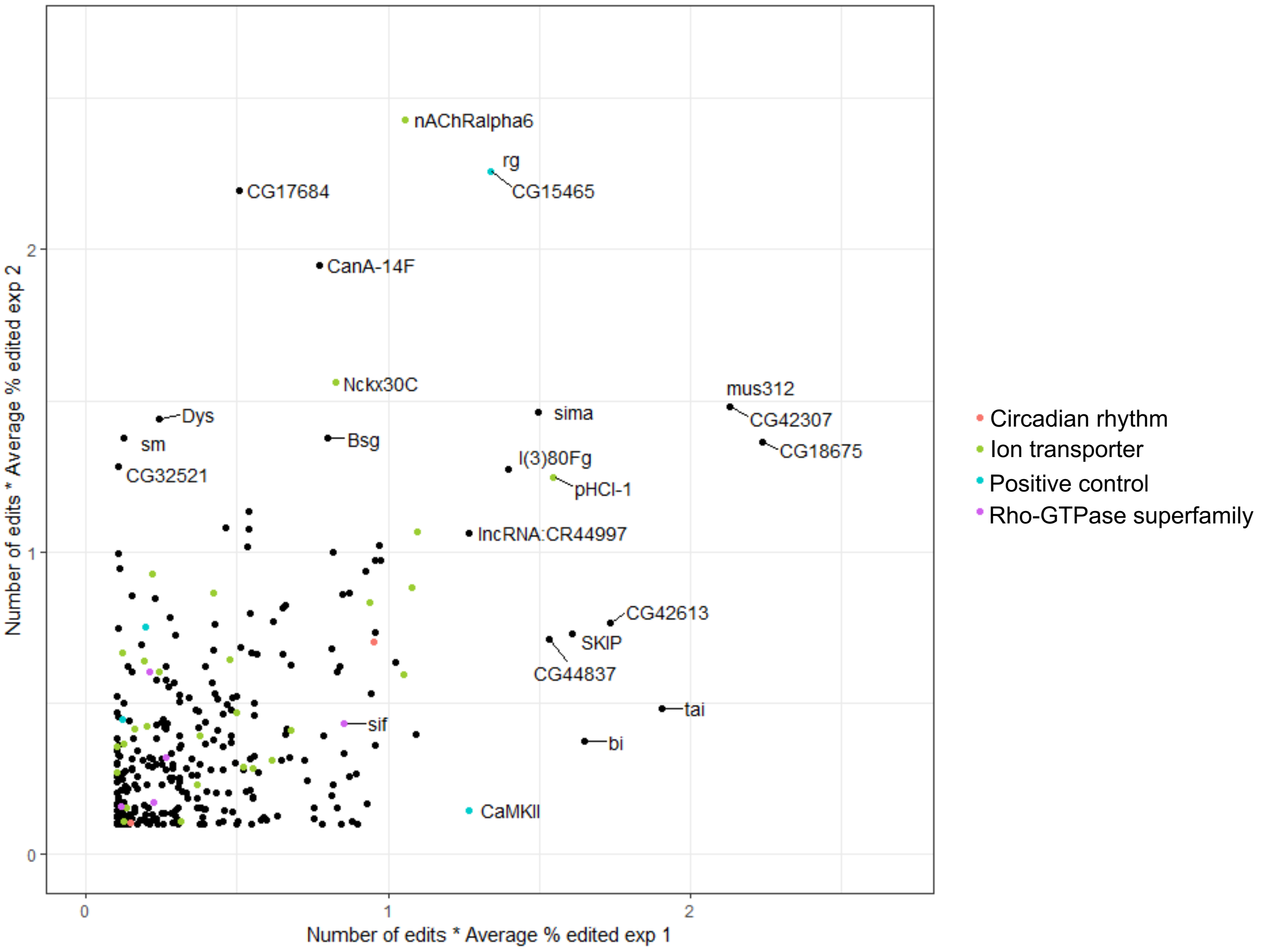


Gundermann et al., Figure S1

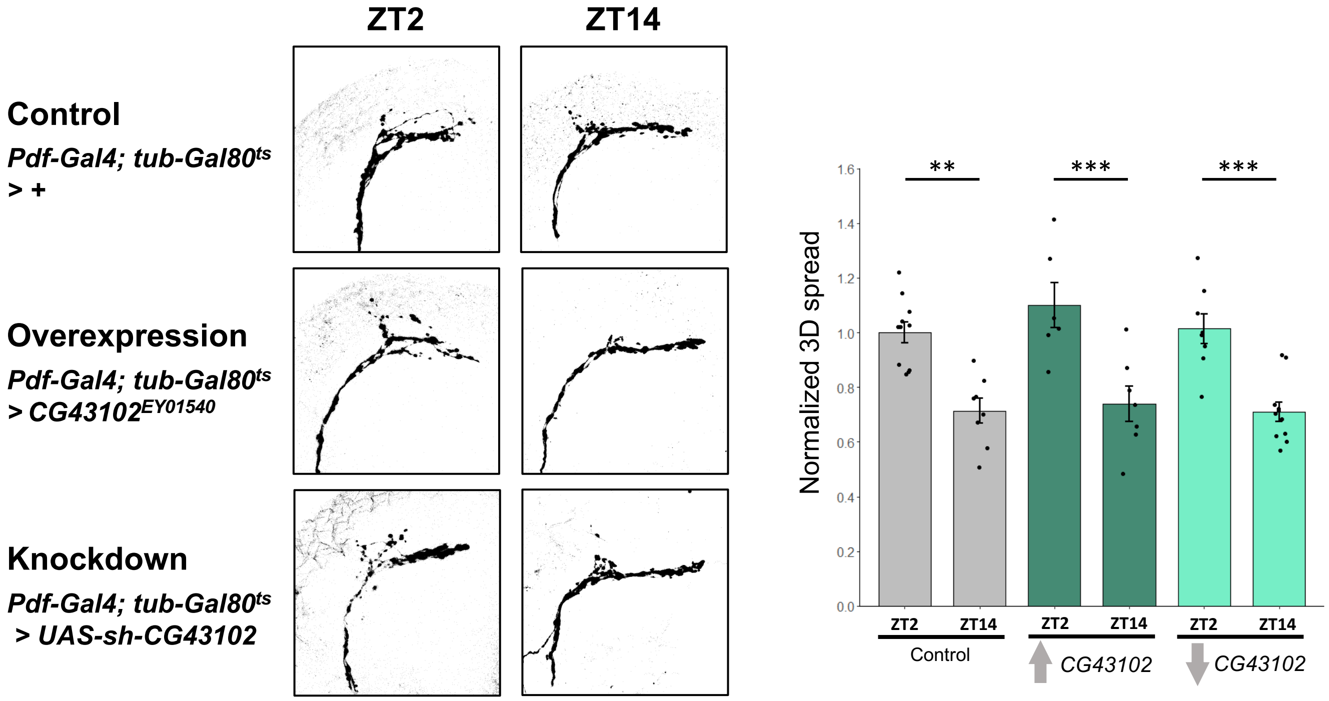
ZT2
ZT14
Control
Pdf-Gal4; tub-Gal80ts
> +
Overexpression
Pdf-Gal4; tub-Gal80ts
> CG43102EY01540
Knockdown
Pdf-Gal4; tub-Gal80ts
> UAS-sh-CG43102
Normalized 3D spread
**
***
***
1.6
1.4
1.2
1.0
0.8
0.6
0.4
0.2
0.0
ZT2
ZT14
ZT2
ZT14
ZT2
ZT14
Control
CG43102
CG43102
